# An Accessible Python Framework for Real-Time Magnetic Tweezers Microscope Control and Image Processing

**DOI:** 10.1101/2025.10.31.685671

**Authors:** James A. London, Abhishek K. Singh, Teague C. Svendsen, Naciye Esma Tirtom, Zachary A. Root, Richard Fishel

## Abstract

Magnetic tweezers are a popular biophysical instrument for manipulating and measuring single molecules. Most groups rely on custom-built setups tailored to specific experiments, making it challenging to implement and share software. Typically, image acquisition and hardware control are automated via LabVIEW, while real-time video processing is implemented in C++/CUDA libraries. Live processing can eliminate the need to store raw video, enabling high throughput, fast acquisition rates, and simplified experimental workflows. However, no open-source general-purpose software framework currently unifies these capabilities for magnetic tweezers experiments. Here, we introduce MagTrack and MagScope open-source Python-based tools designed to fill this gap. MagTrack is an image-processing library that efficiently determines bead positions from magnetic-tweezers videos using CPU or GPU computation. MagScope is a comprehensive software framework offering a graphical user interface, real-time hardware control, data acquisition, and video processing. It is built on a multiprocessing architecture for responsive, high-throughput computation. Together, MagTrack and MagScope offer a fully customizable, end-to-end, open-source Python alternative to proprietary or fragmented systems, enabling laboratories to adapt and extend the framework according to their experimental needs.

## Introduction

Single-molecule force spectroscopy enables direct measurement and manipulation of the mechanics and kinetics of individual biomolecules.^1–3^ Among available force-spectroscopy platforms, magnetic tweezers are popular due to their relative simplicity, low cost, and inherent high throughput.^4–14^ Although these instruments can be built in a variety of configurations, their core concept remains simple: a microscope with a pair of magnets positioned above the sample. In a typical experiment, DNA, protein, or other polymers are tethered between a microscope slide and micron-sized magnetic beads. The magnets exert a tunable force and torque on the beads, controlled by moving the magnets (**Fig. 1a**). With a modest investment in a suitable camera, a capable computer, a few machined components, and standard optical parts, laboratories can construct a high-resolution magnetic tweezers instrument capable of detecting nanometer-scale changes in molecular extension.^15,16^ Beyond their high resolution, magnetic tweezers inherently support parallel measurements, as multiple beads can be tracked simultaneously within a single camera field of view. This high-throughput capability makes magnetic tweezers particularly valuable for studying biological processes that exhibit stochastic behavior, such as enzymatic activity and protein folding.^16,17^

**Figure 1.**
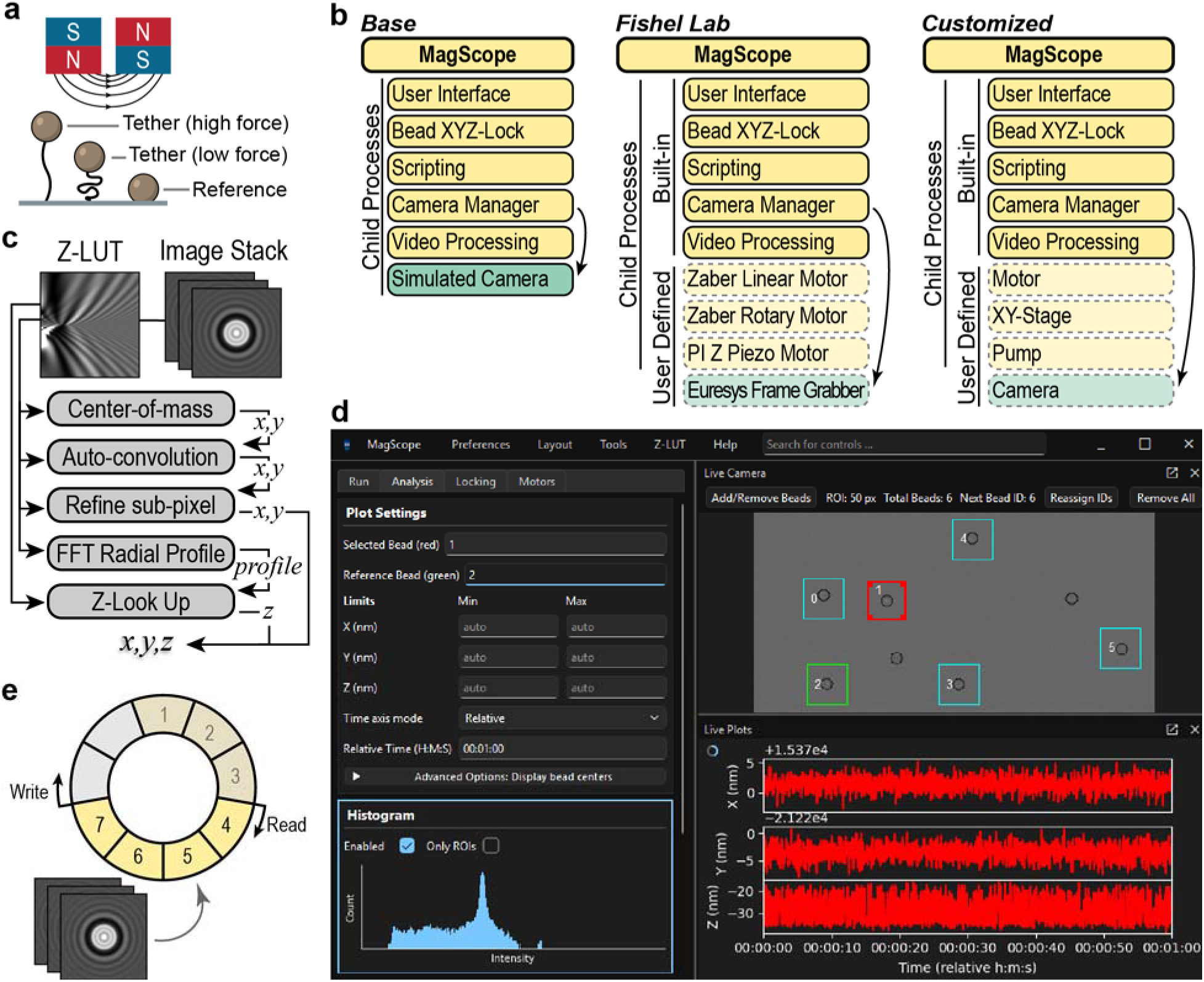
Overview of MagTrack and MagScope. **a** Illustration of magnetic-tweezers principle. **b** Three different use cases of the MagScope framework. *Left)* the “Base” version which requires no additional setup to run. This is used for demonstration purposes and development testing. *Middle)* the “Fishel Lab” implementation which was used to connect our specific hardware and preferred controls to MagScope. *Right)* a “Customized” version to demonstrate the versatility of the MagScope framework for a laboratory-specific implementation. **c** Overview of tracking pipelines that can be composed with MagTrack. **d** Screenshot of MagScope base user interface. **e** Diagram of ring buffers management to provide efficient inter-process access to large datasets including the video stream.

A challenging aspect of building a high-throughput magnetic tweezers system is the software required to process video recordings. Magnetic tweezers videos can be analyzed with generic standalone video-tracking tools such as ImageJ.^17–19^ However, achieving high spatial and temporal resolution demands cameras that generate large images at high frame rates, resulting in enormous data volumes (1-100 GB/min). Handling such data creates a significant bottleneck in experimental workflows. Instead, many instruments are operated with software to process the video feed in real time. This approach not only reduces data storage but also enables researchers to monitor and respond to the results.

A common platform for live processing in magnetic tweezers experiments is LabVIEW, which is well suited for hardware interfacing and instrument control. An influential implementation from the Dekker group combined LabVIEW with a powerful custom-built C++/CUDA library for real-time image analysis.^20,21^ However, LabVIEW is proprietary, expensive, and not widely used outside of instrumentation, limiting its reach. Other C++ and Python implementations have achieved strong performance, but are tailored to individual instruments and lack the documentation, modularity, or general-purpose design necessary for broad adoption.^4,12,16,20–24^

These challenges have highlighted a need for an open-source, accessible, and extensible software framework for magnetic tweezers.^6^ Python is well suited for this goal because it is free, widely used by the biophysics community, supported by mature scientific libraries, and already common in post-processing workflows for magnetic-tweezers data.^25^ A Python-based platform could therefore unify real-time acquisition, image analysis, hardware control, and downstream analysis within a single, modifiable ecosystem.

Here, we present MagTrack and MagScope, complementary open-source Python tools for magnetic-tweezers image processing and real-time microscope control. MagTrack provides CPU- and GPU-compatible bead-tracking algorithms, while MagScope provides a multiprocessing graphical interface for acquisition, hardware coordination, and live analysis. The packages are supported by installation guides, documentation, configuration examples, and benchmarking workflows, enabling other laboratories to adopt, benchmark, and customize the framework for their own instruments. We validate the framework using standard magnetic-tweezers workflows and quantify the tradeoff between complete-pipeline real-time throughput and fixed-bead spatial resolution. The platform supports uninterrupted multi-day acquisition, tracking of thousands of beads at low magnification (10×), and sub-nanometer resolution at high magnification (100×). These results establish MagTrack and MagScope as an accessible, end-to-end framework for magnetic-tweezers experiments.

## Results

### Software framework design

We developed MagTrack and MagScope as complementary open-source Python packages for magnetic-tweezers experiments. MagTrack is a library of bead-tracking algorithms for converting images into x-y-z positions. MagScope integrates these routines into an end-to-end application for live acquisition, hardware control, visualization, scripting, and data storage (**Fig. 1b**). Together, the packages address three practical requirements for adoption: accessibility, modularity, and performance.

To make the framework accessible, we implemented both packages entirely in Python, a free language that is widely used by biophysics groups, supported by mature scientific libraries, and readable enough for users to inspect and modify. Both packages are easily accessed through the Python Package Index (PyPI), maintained on GitHub under the GPL-3.0 open-source license, and run on Windows, macOS and Linux operating systems. To help users get started we have also created installation guides, user guides, and example workflows. To support collaborative development, each package includes testing and benchmarking tools. MagTrack includes a bead-image and Z-LUT simulator to standardize testing and allow new users to try the package without preparing data. Similarly, MagScope starts by default with a simulated camera, allowing users to explore the application before connecting hardware.

Because magnetic-tweezers laboratories use different tracking strategies depending on microscope configuration and experimental goals^12–14,16,20–22,26^, we made the packages modular to support different instrument designs and analysis preferences. MagTrack achieves this with interchangeable methods rather than a single prescribed pipeline. Users can run the default tracking pipeline, combine built-in x-y and z-tracking steps, or add their own algorithms (**Fig. 1c**). MagTrack uses NumPy^27^ and SciPy^28^ for CPU processing and CuPy^29^ for optional GPU processing, allowing one Python implementation of each tracking algorithm to run on either CPU or compatible NVIDIA GPUs. MagScope extends this modular design through templates and base classes for adding hardware interfaces, child processes, and laboratory-specific functionality. Because hardware requirements vary among laboratories, full microscope operation requires manufacturer-provided drivers and implementation of device-specific interfaces.

Finally, we designed MagScope for live performance. It separates camera acquisition, image processing, hardware communication, and the user interface into independent processes. The interface displays the live camera view, selected bead regions of interest (ROIs), and x-y-z trajectories (**Fig. 1d**). Large video and tracking arrays are shared through ring buffers to keep data synchronized while avoiding repeated copying between processes (**Fig. 1e**). MagScope also includes automatic bead detection, Z-LUT generation, ROI recentering (XY-Lock), focus feedback (Z-Lock), optional image storage, and scripting for automated experimental steps. Together, these design choices provide a powerful customizable software foundation for magnetic-tweezers acquisition and analysis.

### Benchmarking throughput and resolution

To evaluate MagTrack performance, we benchmarked the runtime of individual tracking algorithms on synthetic bead-image simulations. We varied ROI size and batch size and calculated throughput as the number of images processed per second. The MagTrack package includes the full benchmarking framework and results from multiple computers. Here, we show a representative set of common algorithms in **Supplementary Fig. 1**. CPU and GPU benchmarks showed that larger ROIs substantially reduced throughput. Batch size had a more complex effect, with throughput generally increasing at larger batch sizes until GPU VRAM became limiting. Overall, these benchmarks indicate that the Python-based MagTrack implementation is fast enough for most magnetic-tweezers workflows on our hardware, reaching >10^6^ images/s for 16×16-pixel ROIs and >10^3^ images/s for 256×256-pixel ROIs.

Next, we benchmarked real-time processing of MagScope/MagTrack on a magnetic-tweezers instrument (**Supplementary Fig. 2**). We measured the tradeoff between throughput and resolution by varying objective magnification, camera binning, effective pixel size, and ROI size while tracking fixed 3-μm-diameter reference beads at 100 Hz (**Fig. 2a**). For each configuration, we determined the maximum number of bead ROIs that could be tracked continuously without dropped frames (**Fig. 2b,c**). In the highest-resolution condition tested, 50 nm/pixel sampling with 256×256-pixel ROIs, the system tracked 9 beads simultaneously. In the lowest-resolution condition tested, 1000 nm/pixel sampling with 16×16-pixel ROIs, the system tracked 3,448 beads simultaneously. At a higher acquisition rate of 1000 Hz, a high-throughput configuration using 10× magnification, 1000 nm/pixel sampling, and 16×16-pixel ROIs supported continuous tracking of 237 beads (**Fig. 2c**; **Supplementary Fig. 3**).

**Figure 2.**
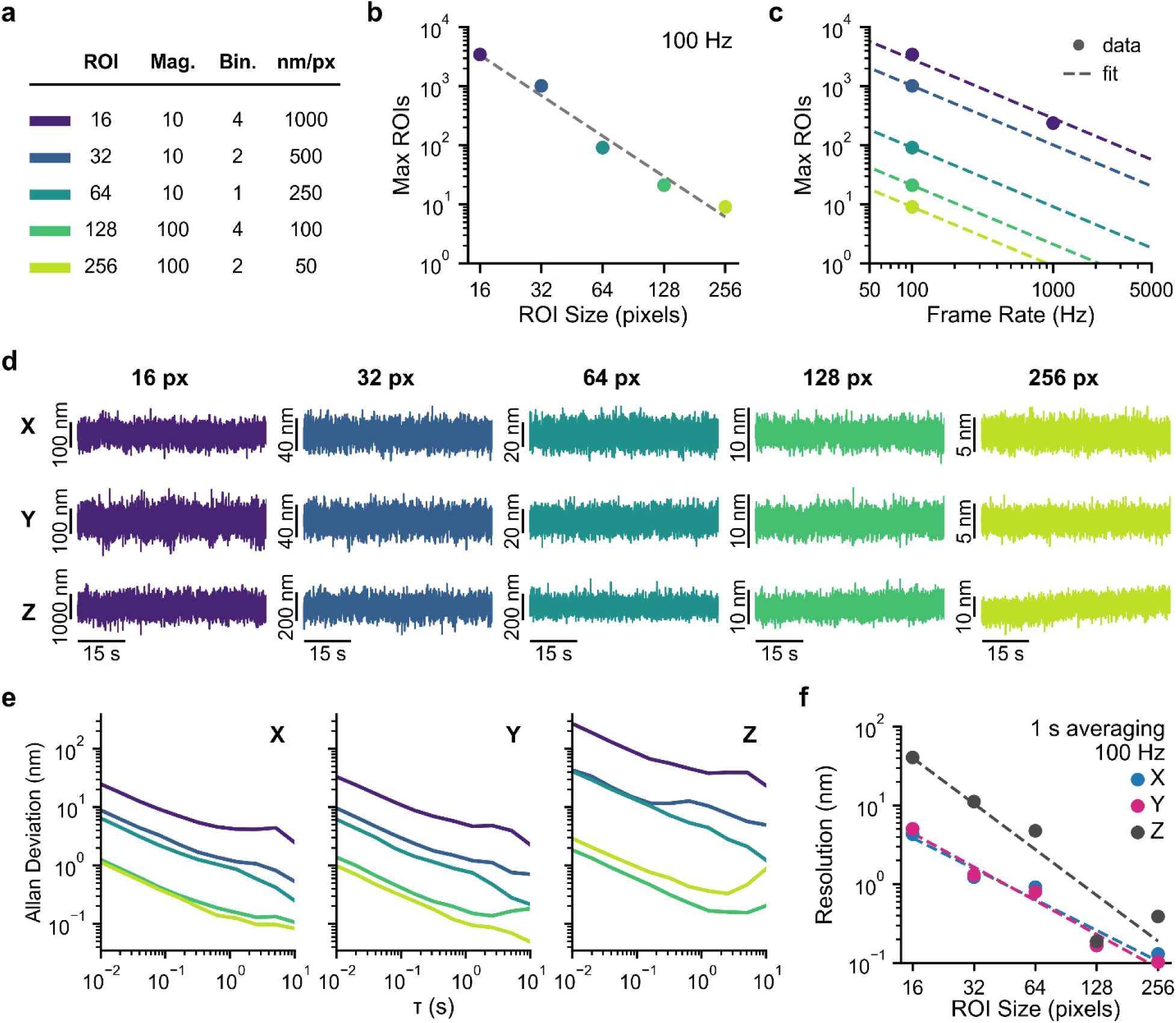
Throughput-resolution tradeoff of the complete MagScope/MagTrack pipeline. **a** Benchmark imaging configurations, showing ROI side length (“ROI”), objective magnification (“Mag.”), camera pixel binning (“Bin.”), and final pixel resolution (“nm/px”); colors are used consistently across panels. **b** Maximum number of bead ROIs tracked continuously at 100 Hz without dropped frames as a function of ROI size on the tested hardware. Points show measured values and the dashed line shows the empirical fit. **c** Maximum supported bead ROIs as a function of acquisition rate for each imaging configuration. Points show measured values and dashed lines show fitted trends. **d** Representative reference-subtracted x-y-z traces from 3-μm-diameter fixed beads for each ROI size, showing increased positional noise with coarser sampling. **e** Allan deviation of the traces in panel d for x-y-z coordinates as a function of averaging time, τ. **f** Spatial resolution after 1 s averaging at 100 Hz for x-y-z tracking across ROI sizes. Together, these measurements define the practical tradeoff between high-throughput acquisition and high-resolution bead tracking for MagScope/MagTrack on the tested hardware.

To define the resolution limit of the high-resolution configuration, we next analyzed fixed beads at 100× magnification, 50 nm/pixel sampling, and 256×256-pixel ROIs. Fixed 3-μm-diameter reference beads were tracked at 145 Hz with Z-Lock feedback enabled, and Allan-deviation analysis gave maximum resolutions of 0.29, 0.25, and 0.55 nm in x-y-z (**Supplementary Fig. 4a**). These values include shared instrument drift and low-frequency vibration across the field of view. Subtracting the position of one fixed reference bead from another removed this shared motion and improved the maximum resolution to 0.04, 0.03, and 0.13 nm in x-y-z (**Supplementary Fig. 4b**).

We then used this reference-subtracted fixed-bead metric to compare tracking precision across the same ROI sizes and effective pixel sizes used for the throughput benchmark. Representative reference-subtracted x-y-z traces and Allan deviations showed that larger ROIs and finer effective pixel sizes improved spatial resolution, whereas smaller ROIs and lower magnification supported higher multiplexing (**Fig. 2d,e**). The resolution was after 1 s averaging improved with larger ROIs across x, y, and z (**Fig. 2f**). Together, these measurements demonstrate the practical operating range of MagScope/MagTrack on our tested computer, camera, and frame-grabber hardware, and show that the complete Python pipeline supports both high-resolution and high-throughput magnetic-tweezers acquisition.

### Long-term stability with XY-Lock and Z-Lock

For extended measurements, MagScope includes optional XY-Lock and Z-Lock routines that maintain bead ROIs and focus during acquisition. XY-Lock periodically recenters ROIs to keep beads within the tracked window, but bead positions are reported in camera field-of-view coordinates rather than ROI-local coordinates. As a result, ROI recentering does not redefine the reported bead position. Z-Lock uses the measured z position of a selected reference bead to adjust the focus piezo and keep the field of view within the calibrated Z-LUT range.

We tested whether these routines introduced discontinuities or measurable excess noise by tracking the same fixed reference beads for 2 h at 100 Hz with XY-Lock and Z-Lock enabled or disabled. During this period, uncompensated x-y drift required approximately 3 to 6 pixels of ROI correction. With XY-Lock enabled, one-pixel ROI updates kept beads centered without introducing discontinuities into the reported x-y traces or the differential trace between reference beads (**Supplementary Fig. 5a,b**). Additionally, Z-Lock stabilized the z position of the selected reference bead without measurably changing the Allan deviation of the differential z trace (**Supplementary Fig. 5b**). Using the same high-resolution configuration, MagScope tracked reference beads continuously for more than 4 days (**Supplementary Fig. 4d**). These measurements show that MagScope can maintain stable real-time tracking during extended acquisition while preserving x-y-z position measurements for downstream analysis.

### Tethered DNA measurements and hairpin kinetics

After defining fixed-bead resolution and acquisition stability, we next asked whether reference-subtracted tracking behaved as expected for DNA-tethered beads, where Brownian motion rather than instrumental noise dominates the fluctuations. Magnetic beads with a 2.8-μm diameter were tethered to the flowcell by 4 kb DNA molecules and held at 1 pN of force. Tethered beads and surface-fixed reference beads were tracked at 100 Hz with Z-Lock enabled, and reference-subtracted traces were used to suppress shared drift while preserving Brownian fluctuations of the tethered bead (**Supplementary Fig. 4c**). The corresponding Allan deviations were higher than those of fixed reference beads, as expected for tethered beads at low force, and followed the trend predicted by a damped harmonic oscillator model^30^ (**Supplementary Fig. 6**).

We then tested whether MagScope could combine this tethered-bead tracking with scripted hardware control during a standard magnetic-tweezers kinetics measurement. We used a 1 kb DNA construct containing a nicked 60 nt palindromic hairpin (**Fig. 3a**). Beads were tracked at 145 Hz in the high-resolution configuration described above, and MagScope scripting was used to perform force ramps from 6 to 20 pN at 0.2 pN/s.

**Figure 3.**
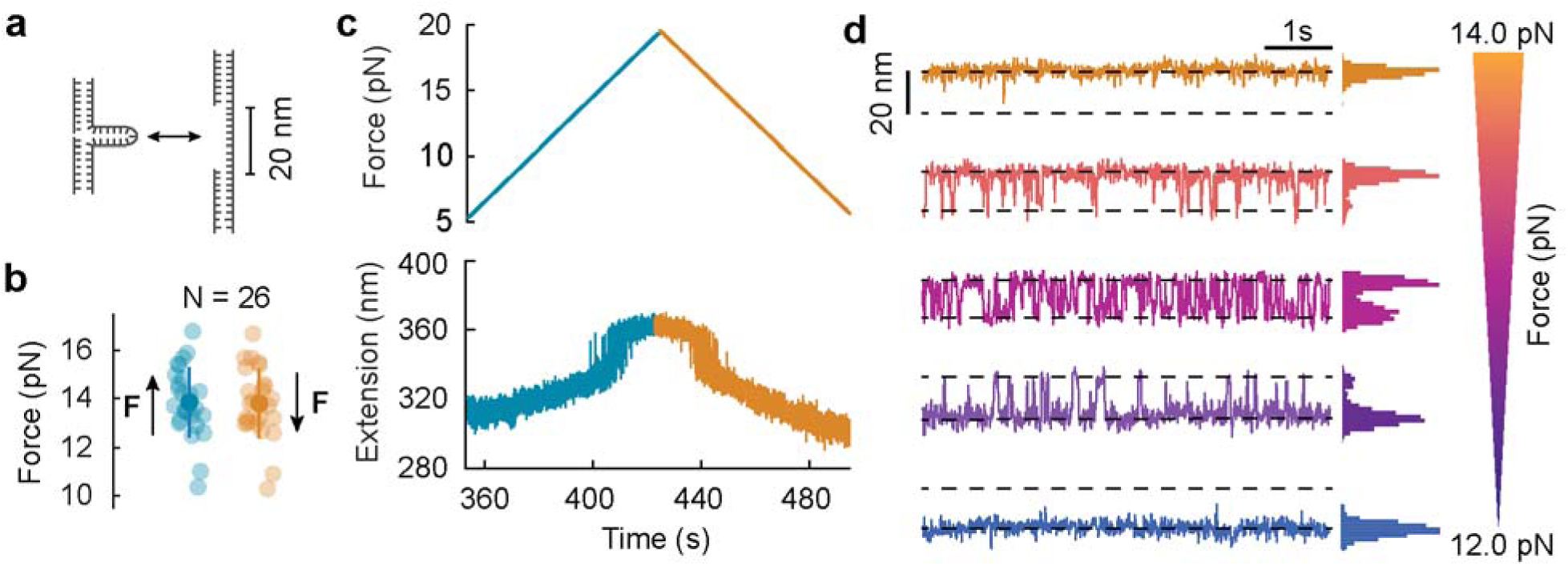
Dynamics of DNA hairpin folding. **a** Schematic of a reversible two-state DNA hairpin. **b** Distributions of rupture and reformation forces of 26 molecules with mean (opaque circle) and standard deviation (error bar) are shown. Rupture forces were measured during increasing-force ramps, and reformation forces were measured during decreasing-force ramps. **c** Force ramp from 6 pN to 20 pN to 6 pN at 0.2 pN/s and its response on a hairpin DNA is shown by a representative trace. The hairpin ruptures as the force increased at ∼410 s and reforms as the force decreased at ∼440 s. **d** Two-state kinetics of nicked hairpin DNA at discrete equilibrium forces from 12 pN to 14 pN at steps of 0.5 pN shown as occupancy histograms with kernel-density estimates. The population shifts from predominantly open to predominantly closed hairpin state with decreasing force, with an even occupancy near 13 pN. All measurements were performed at 145 Hz in a high-resolution configuration using 2.8-μm-diameter M-280 as tethered beads (100× magnification; 50 nm/pixel; 256×256-pixel ROIs).

During force ramps, single-molecule extension traces resolved discrete 20 nm jumps corresponding to hairpin rupture as force increased and hairpin reformation as force decreased (**Fig. 3c**). Across 26 molecules, rupture and reformation forces were centered near 13.7 pN (**Fig. 3b**). In force-clamp measurements from 12 to 14 pN, the hairpin repeatedly opened and closed, producing two-state extension traces whose open-state occupancy increased with force (**Fig. 3d**). These measurements demonstrate that MagScope can coordinate force control, scripted experimental protocols, and real-time x-y-z tracking to resolve nanometer-scale conformational kinetics.

### High-throughput flow stretching

Finally, we tested a practical high-throughput experiment with MagScope/MagTrack using a flow-stretching assay on a 4 kb DNA construct^14,17,31^. Experiments were performed at 10× magnification with 500 nm/pixel sampling and 28×28-pixel ROIs (**Fig. 4a**). Under these conditions, MagScope simultaneously tracked 1,000 1-μm-diameter beads at 100 Hz (**Fig. 4b**). The full set of raw trajectories, a trajectory-density map, and a representative field of view is shown in **Supplementary Fig. 7**.

**Figure 4.**
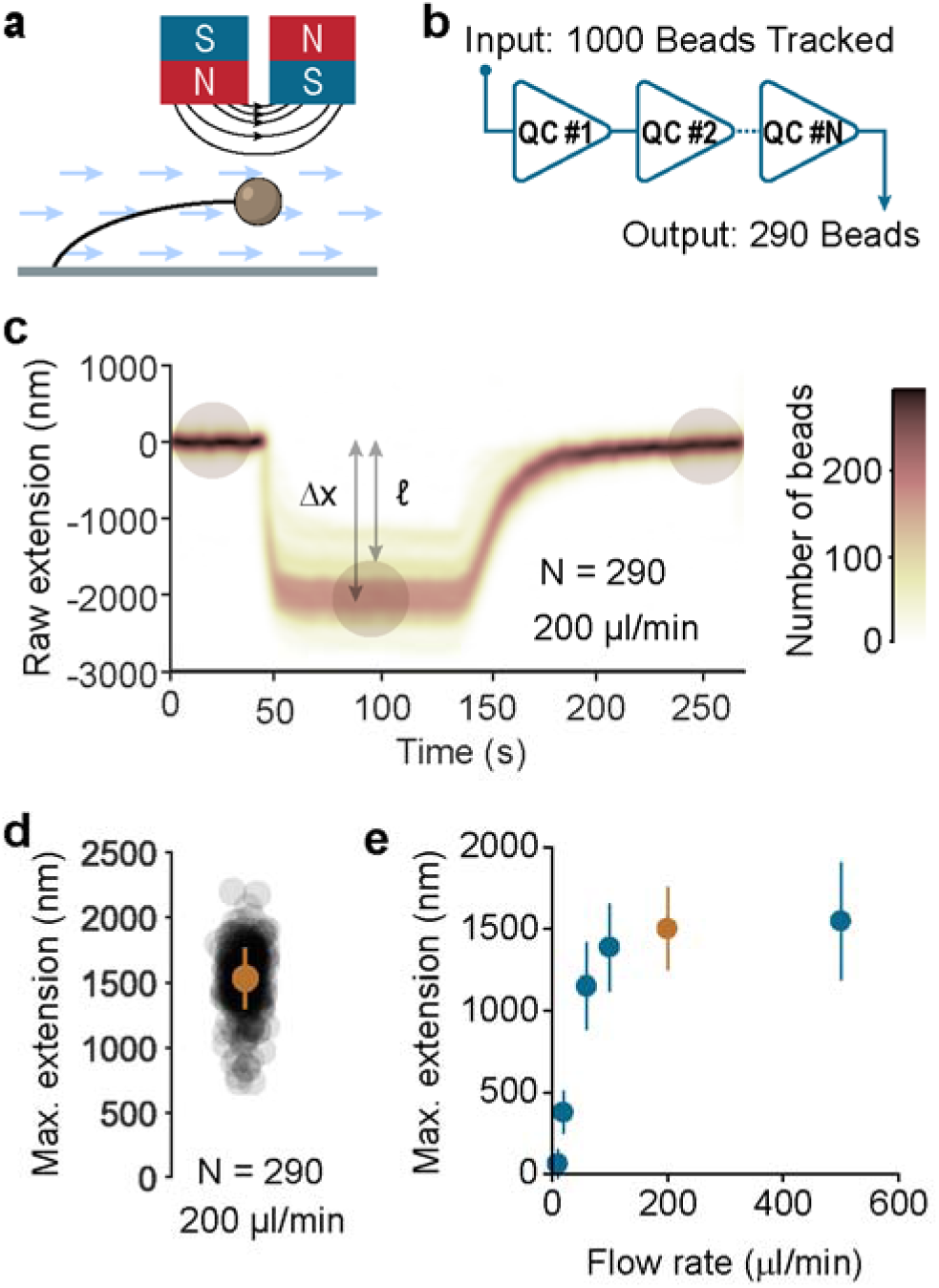
Flow stretching 1,000 molecules. **a** Illustration of the flow stretching configuration: a 4 kb DNA tethered at one end to a glass surface and at the other end to a 1-μm magnetic bead. A constant magnetic force (1 pN) and flow-rate dependent hydrodynamic drag force is applied to the magnetic bead. **b** Schematic of the data processing scheme. 1,000 beads were simultaneously tracked, while registering their x-y-z positions. These tracks were then processed through a series of quality-control filters to eliminate beads showing atypical trajectories. **c** The 2D kernel-density of real-time raw bead positions (from bead-center displacement) during a step of steady flow of 200 μL/min, followed by subsequent relaxation upon return to zero flow is shown. The kernel-density was mapped for the accepted 290 molecules, selected from 1,000 initially tracked. **d** Distribution of maximum DNA extension, *ℓ*, for the QC-accepted subset (N = 290) after correcting raw extensions, Δx, shown in panel c by subtracting the bead radius (500 nm). Marker shows mean and standard deviation. **e** Maximum DNA extension at different flow rates is shown. Extension approaches the 4 kb contour length (∼1.37 µm) by ∼200 µL/min with no further extension observed at higher rates. All measurements were made at 100 Hz in a high-throughput configuration using 1-μm-diameter C1 beads (10× magnification; 500 nm/pixel; 28×28-pixel ROIs).

A constant magnetic force of 1 pN was applied throughout the experiment, and buffer flow through the flowcell stretched the DNA tethers. DNA extension increased during flow and returned toward the initial value when flow stopped (**Fig. 4c**). Quality-control filters removed trajectories consistent with stuck beads, multiple tethers, broken tethers, or failure to return to baseline, yielding 290 high-quality traces for quantitative analysis (**Fig. 4b,c**). This filtering reflects experimental yield and tether quality rather than the number of beads tracked by the software.

Across flow rates, DNA extension increased with flow and plateaued near the expected 1.36 μm contour length of the 4 kb DNA construct by 200 μL/min (**Fig. 4d,e**). These results demonstrate that MagScope and MagTrack can perform real-time high-throughput magnetic-tweezers acquisition while retaining enough trajectory-level information for quality control and downstream molecular analysis.

## Discussion

Here we describe MagTrack and MagScope, complementary open-source Python packages for magnetic-tweezers image processing and real-time microscope control. Because magnetic-tweezers instruments are commonly custom-built, complete software workflows must often be customized for specific hardware configurations and experiment-specific workflows. This customization can make otherwise capable instruments difficult to reproduce, benchmark, or adapt outside the laboratory in which they were developed. MagTrack and MagScope provide an alternative to LabVIEW-based acquisition and C++/CUDA-based tracking workflows by implementing these functions in an extensible Python framework.^20,21^ The packages were designed around three practical requirements for adoption: accessibility, modularity, and performance. In practice, this means that users can install and test the software without first connecting a complete instrument, replace or extend individual tracking and hardware-control components, and benchmark the complete acquisition pipeline under their own experimental conditions.

To evaluate the framework, we performed end-to-end benchmarking during live acquisition on a magnetic-tweezers instrument. We varied objective magnification, camera binning, effective pixel size, and ROI size, then measured both fixed-bead precision and the maximum number of bead ROIs that could be tracked continuously without dropped frames. These measurements showed the expected tradeoff between spatial resolution and the number of beads tracked. Higher magnification, finer effective sampling, and larger ROIs improved x-y-z precision, whereas lower magnification and smaller ROIs increased the number of bead ROIs that could be tracked at 100 Hz or 1000 Hz. In the highest-resolution condition tested, the MagScope/MagTrack pipeline achieved sub-nanometer reference-subtracted x-y-z resolution. In the highest-throughput condition tested at 100 Hz, the system tracked 3,448 bead ROIs continuously without dropped frames, while at 1000 Hz it supported continuous tracking of 237 bead ROIs. These results demonstrate that our framework can support magnetic-tweezers experiments ranging from high-resolution bead tracking to high-throughput acquisition, depending on their spatial, temporal, and throughput requirements.

Beyond fixed-bead benchmarking, the hairpin and flow-stretching experiments show that MagTrack and MagScope can support complete magnetic-tweezers workflows. In the hairpin assay, MagScope combined real-time x-y-z tracking with scripted magnet control, force ramps, force clamps, and tether validation. This allowed us to resolve reversible 20 nm hairpin rupture and reformation events during force ramps, and to measure force-dependent two-state kinetics under constant force. In the flow-stretching assay, the same framework was configured for high-throughput acquisition, tracking 1,000 beads at 100 Hz while retaining trajectory-level information for downstream filtering and analysis. Quality-control filtering recovered 290 high-quality single-tether traces, and the accepted traces showed flow-dependent extension of the 4 kb DNA construct. Together, these experiments show that MagTrack and MagScope can be used for both detailed single-molecule kinetics and high-throughput trajectory collection, producing datasets consistent with the expected behavior of standard magnetic-tweezers assays.

These validation experiments also clarify the scope of the contribution relative to previous magnetic-tweezers software. Prior implementations established that real-time tracking of spherical beads can be achieved with optimized acquisition programs and dedicated image-processing code.^12,16,20–22^ MagTrack and MagScope address the complementary need for a framework in which tracking methods, hardware interfaces, acquisition logic, scripted protocols, and benchmarking can be modified within an open-source environment. This distinction is important because laboratories often need to preserve instrument-specific control while still using software that can be inspected, tested, and extended by other users. The goal is not to prescribe one optimal magnetic-tweezers software implementation, but to provide a common structure in which different implementations can be built, compared, and maintained.

Several limitations should be considered when adapting MagTrack and MagScope to other instruments. First, complete microscope control requires device-specific interfaces, because cameras, frame grabbers, motors, piezo controllers, pumps, and other components use different drivers and communication protocols. MagScope provides templates and base classes for these interfaces, but laboratories will still need to implement and validate the components specific to their hardware. Second, GPU acceleration requires compatible NVIDIA hardware, CUDA-compatible Python packages, and matching driver and library versions. CPU-based tracking remains available, but the achievable throughput will depend on the selected tracking routine, ROI size, bead number, and frame rate. Finally, the throughput and resolution values reported here are specific to the camera, frame grabber, computer, optical configuration, bead size, and imaging conditions used in this work. For this reason, end-to-end benchmarking should be performed when the framework is deployed on a new instrument. Within these limits, MagTrack and MagScope provide a practical foundation for reproducible, extensible, and community-adaptable magnetic-tweezers acquisition. By combining open-source implementation with modular hardware integration and complete-pipeline benchmarking, the framework can be evaluated and modified as experimental requirements change.

## Methods

### Image processing algorithms

Center-of-mass: This is a simple routine to obtain initial estimates of bead positions from square image stacks. The input image stack should be arranged as a three-dimensional array of size N×N×T, where N is the number of pixels and T is the number of frames. For each frame, the algorithm treats the pixel intensities as a mass distribution, and computes x and y centroid coordinates. Before calculating the centroids, each frame can optionally be background-corrected by subtracting either the frame-wise mean or median intensity and taking the absolute value, which reduces the influence of uniform background offsets. The corrected images then collapse along the x- and y-axes to obtain one-dimensional intensity profiles, and the first moment of these profiles is evaluated with respect to the discrete pixel coordinate grid. Dividing these moments by the total frame intensity yields the x and y positions for each frame.

Auto-convolution: This is used to refine bead center positions given an initial estimate. For each frame in the image stack, given x and y coordinates were used to select the corresponding central column and row. These one-dimensional intensity profiles are mean-subtracted and multiplied by a Gaussian window centered on the estimated position, which suppresses edge contributions and emphasizes the region near the bead. The algorithm then computed the auto-convolution of each profile using an FFT-based convolution and identified the location of the maximum in the row and column convolutions. Because the auto-convolution peak is centered on the point of highest symmetry, these peak positions were converted back to image coordinates to obtain updated x and y bead positions for each frame. When needed for later sub-pixel fitting, the full convolution profiles can be retained instead of only the peak indices.

Sub-pixel auto-convolution: This feature is identical to auto-convolution. Once the convolution and maximum are found, a small, fixed-width neighborhood (typically 5 points) is extracted and fit with a quadratic function. The vertex of this fitted parabola provides a sub-pixel estimate of the convolution peak position along the row and column. These refined peak locations are then mapped back into the original image coordinate system, accounting for the symmetry of the auto-convolution, to yield sub-pixel x and y bead coordinates for each frame.

Auto-convolution multiline: This is the same as the auto-convolution but rather than one central row/column being evaluated, multiple rows/columns are averaged. The number of rows/columns can be provided by the user.

Sub-pixel auto-convolution multiline: The same as the auto-convolution multiline routine but with the sub-pixel fitting from the sub-pixel auto-convolution routine.

QI (Quadrant Interpolation): Generally, this is the same as the originally developed approach.^20^ For each frame, the algorithm samples the intensity along the horizontal and vertical axes that pass through the current bead position, extracting short one-dimensional profiles centered on the estimated pixel. For each axis, three neighboring samples (center pixel and its immediate neighbors) are used to fit a quadratic function to the local intensity profile. The vertex of this fitted parabola provides a sub-pixel offset relative to the central pixel along that axis. These x- and y-offsets are then added to the input coordinates to yield refined bead positions for each frame.

Radial Profile: This is used to convert each bead image into a one-dimensional intensity profile as a function of distance from the bead center. For each frame in the image stack, the routine takes the corresponding x and y center coordinates and constructs a grid of pixel positions. It then computes the Euclidean distance from the center to every pixel and assigns each pixel to a radial bin based on this distance. An optional oversampling factor subdivides the native pixel-wide bins into finer radial intervals, increasing the resolution of the profile without changing the underlying image. Within each radial bin, the pixel intensities are averaged, yielding a smooth radial intensity profile for each frame.

FFT Radial Profile: For each frame in the stack, we computed a 2D real-valued only discrete Fourier transform and shifted the zero-frequency component to the center of the spectrum. The complex Fourier coefficients were converted to their magnitudes and then azimuthally averaged by grouping coefficients according to their radial distance from the origin in frequency space. To increase radial resolution without changing the underlying data, the radial coordinate was oversampled by an integer factor, and the magnitudes were accumulated into these oversampled radial bins. The resulting one-dimensional profiles were restricted to a user-defined range. Because this procedure operates directly on the Fourier magnitude, it does not require prior knowledge of the bead center.

Z-Look-up: We used a look-up-table-based routine to convert radial intensity profiles into z positions. For each frame, the bead image was first reduced to a one-dimensional radial profile as described above. These profiles were then compared to a precomputed Z-look-up table (Z-LUT) containing reference radial profiles acquired at known z positions. In the Z-LUT, the first row stores the calibrated z coordinates, and the remaining rows store the corresponding reference profiles (with the central pixel excluded from matching). For each measured profile, the routine computes the Pearson correlation coefficient against every reference profile in the Z-LUT and identifies the reference index that yields the highest correlation. To obtain sub-plane accuracy, the correlation values in a small neighborhood around this maximum are fitted with a parabola, and the vertex of this fit is used to estimate a fractional index between LUT entries. This fractional index is then interpolated onto the calibrated z-axis stored in the first row of the Z-LUT, yielding a continuous estimate of the bead’s z position for each frame.

Standard Pipeline: First the center-of-mass is called, the results are passed to auto-convolution, the results are then updated with sub-pixel auto-convolution multiline five times, then the radial profile is calculated, and finally the Z-look-up if a Z-LUT is provided.

### XY-Lock and Z-Lock

The XY-Lock works by checking the most recent valid position of all currently selected beads either when the user requests it with a button push or when the user has enabled a timer. The timer can be set to any interval the user requests; we used ten seconds. The difference between the bead’s positions and the center of the ROI is then calculated for both the x and y directions separately. Differences of less than one pixel for either direction are ignored. Larger differences are rounded to the nearest pixel and then the ROIs are moved. An upper limit can be specified by the user to prevent sudden large changes. Overall, this routine does not disrupt the bead’s measured position because the positions are reported as the distance from the top left corner of the camera to the center of the bead. Thus, the position of a bead is the sum of the bead’s position relative to the ROI and the ROI relative to the camera’s field-of-view. If the ROI moves one pixel to the left, the bead will appear to move one pixel to the right relative to the ROI, so no change is observed. However, if the bead is too close to the edge of the ROI, a more noticeable change can be apparent due to the improved performance of the algorithms when the bead is near the center of the ROI.

Z-Lock can also be triggered manually or on a timer. Rather than updating all bead ROIs, it uses the measured z position of one user-selected reference bead. The user specifies a target z position, and MagScope calculates the difference between the current and target z values. This offset is then added to the objective focus-piezo target position.

### Microscope design, computer and software environment

We built an inverted microscope following previously established magnetic-tweezers designs, and an image of the assembled instrument is shown in **Supplementary Fig. 2**^11,15,16,20,32,33^. An objective lens (UPlanSApo 100×/1.40 oil-immersion or UPlanSApo 10×/0.40 NA, Olympus, Japan) was mounted on a single-axis piezo actuator (P-721.CDQ, Physik Instrumente) rigidly integrated into a metal microscope frame, enabling precise motion along the optical (z) axis relative to a fixed flowcell. A separate optical rail post carries a light source (M530L2, Thorlabs) and a linear actuator (Thorlabs X-LSQ075A-E01) with pulley system. The linear actuator translates the pulley system. One wheel of the pulley is attached to a rotary stepper motor (Zaber X-NMS17-E01) and the other pulley is attached to a 3D printed magnet holder that separates two 0.25 inch cube N52 neodymium magnets (B444-N52, K&J Magnetics) by 0.9 mm. The entire vertical post is mounted on a XY-stage (Thorlabs TBB0606) to position the magnet pair in such a way that the light source, center of magnet pair and the objective lens share a common optical axis. The linear actuator translates the pulley/magnet assembly along the optical axis to vary magnet-flowcell separation. MagScope coordinates all motors, executing programmed protocols for DNA extension and supercoiling. Images were captured using a CMOS camera (Vieworks VC-25MX2-M150I) and transferred to a CoaXPress frame-grabber (Euresys PC3622). The computer used for the magnetic tweezers experiments was equipped with 64 GB of RAM, an Intel i5-12600KF CPU, NVIDIA GeForce RTX 3060 Ti GPU, NVIDIA CUDA Toolkit 13.0, NVIDIA driver 581.95, and Python 3.11.3.

MagScope/MagTrack can be run in CPU mode without GPU-specific dependencies. CPU-based tracking is therefore the default supported configuration. Optional GPU acceleration requires a compatible NVIDIA GPU, appropriate NVIDIA drivers, CUDA-compatible Python packages, and matching CUDA libraries. Because GPU performance depends on the specific hardware and software environment, GPU acceleration should be validated on each user’s system.

MagTrack was tested on Windows, macOS, and Linux with Python 3.8 and later. GPU support with MagTrack is not available on macOS because CUDA-compatible NVIDIA GPUs are not supported on current macOS systems. GPU support was tested with CUDA 12 and CUDA 13 on Windows and Linux. Testing logs and benchmark results are provided in the benchmarking section of the repository. MagScope was tested on Windows, macOS, and Linux with Python 3.10 and later.

### Algorithm specific benchmarking on simulated data

MagTrack includes a benchmarking framework with benchmarking logs included in the repository. Additionally, in this work performance was benchmarked (**Supplementary Fig. 1**) using simulated image stacks containing a single centered bead at z = 0 nm. Five imaging configurations were tested: 16×16, 32×32, 64×64, 128×128, and 256×256-pixel ROIs, corresponding to effective sampling distances of 1000, 500, 250, 100, and 50 nm/pixel, respectively. Images were generated at higher spatial sampling and mean-binned to the final ROI dimensions, then replicated into contiguous double-precision image stacks. These parameters were selected to align with the practical end-to-end benchmarking performed with MagScope. Batch sizes increased in powers of two from 2^4^=16 to 2^18^=262,144 images. ROI/batch combinations requiring more than 2 GB of input memory were excluded so as to not fill the GPU RAM during computation. Benchmarking was performed on the same computer as all other testing; it is equipped with 64 GB of RAM, an Intel i5-12600KF CPU, NVIDIA GeForce RTX 3060 Ti GPU, NVIDIA CUDA Toolkit 13.0, NVIDIA driver 581.95, and Python 3.11.3.

The center-of-mass, sub-pixel auto-convolution, radial-profile, FFT-profile, and z-lookup functions were benchmarked independently on the CPU and GPU. Initial x-y coordinates were set to the center of each ROI. For z-lookup, a simulated Z-LUT spanning −10,000 to 10,000 nm in 200-nm increments was used. Z-LUT generation and radial-profile preparation were performed before timing and were not included in the reported z-lookup runtime. Each condition was preceded by 10 untimed warm-up calls and measured over 10 subsequent calls using Python’s high-resolution time.perf_counter_ns timer. CUDA operations were synchronized immediately before and after each GPU measurement. Image simulation, stack allocation, CPU- to-GPU data transfer, result validation, and memory cleanup were excluded from the timed intervals. Throughput was calculated as the batch size divided by runtime.

### Flowcell assembly

Flowcells were assembled by sandwiching a 0.12 mm-thick double-sided tape spacer between two glass coverslips (each 0.17 mm thick). The original coverslips were each 24 × 50 mm. The bottom coverslip was used at its full size (24 × 50 mm), while the top one was trimmed to 24 × 40 mm. The double-sided tape was cut into a rectangle to define a channel of dimensions 2 × 42 mm and positioned between the coverslips in such a way that ∼1 mm of channel length remained open at each end to serve as the inlet and outlet. A 0.28 mm inner diameter polyethylene tube (BD Intramedic, PE10) was used for the inlet whereas a 0.76 mm inner diameter polyethylene tube (BD Intramedic, PE60) was used for the outlet of the flowcell. The tubes were affixed at each end, using a short section of pipette tip as a coupling, to provide the inlet and outlet connections. The resulting flowcell volume was typically ∼10 µL.

### Experiment procedures

Magnetic tweezers experiments were carried out using a standard procedure^10,15,16,31^. First the flowcell was rigidly mounted and the outlet was connected to a pump (kD Scientific LEGATO, 78-8110) with a 10 ml syringe. The flowcell was then washed with a 400 μl of purge buffer (10 mM Tris pH 7.5; 140 mM NaCl; 1 mM EDTA; 0.5% Tween-20). A solution of 25 μl of polystyrene beads (Duke Standards, 4K-03, 3-μm-diameter) with 75 μl TNE buffer (10 mM Tris pH 7.5; 140 mM NaCl; 1 mM EDTA) was vortexed and injected into the flowcell followed by two hours of incubation to non-specifically fix polystyrene beads onto the flowcell surface. A further wash with TNE buffer removed all the polystyrene beads that were not attached to the surface. These polystyrene beads served as fixed reference beads to correct for mechanical drift during the measurements. The flowcell surface was then treated with 100 μl of 0.1 mg/ml Digoxigenin antibody followed by 2 hours of incubation. Thereafter, the flowcell was passivated using 100 μl of BSA solution, TNEB buffer (10 mM Tris pH 7.5; 140 mM NaCl; 1 mM EDTA; 0.2 mg/ml BSA), with 30 minutes of incubation. A 100 μl solution of 2 pM DNA, having digoxigenin at one end and biotin at the other, was prepared in TNEB and injected followed by 20 minutes of incubation. An additional wash with TNEB removed untethered DNA. Thereafter, streptavidin coated magnetic beads were injected. For the high-resolution experiments in **Supplementary Fig. 4c** and **Fig. 3** 2.8-μm-diameter M-280 streptavidin beads (ThermoFisher, 11205D) were used. For the high-throughput experiments in **Fig. 4**, 1-μm-diameter streptavidin C1 beads (ThermoFisher, 65001) were used. In both cases for each experiment ∼10^6^ beads were first prepared by washing them in 100 μl of TNEB twice; the beads were then injected into the flowcell and excess unbound beads were washed out with TNE buffer.

Additional purge of non-specifically attached magnetic beads was performed by applying 20 pN of force for M-280 beads and 5 pN of force for C1 beads, followed by a TNE wash. While measuring the kinetic response from palindromic hairpin DNA, the tethered DNA was subjected to a series of test ramps. For instance, 20 (+) turns and 20 (−) turns at 1 pN were used to ensure that extension of DNA remains unchanged during this scan. A single tethered DNA showed no change in DNA extension, whereas a multi-tethered DNA showed decrease in extension since the DNAs become entwined.

For the experiments in **Supplementary Fig. 4** and **Fig. 3** imaging was performed with a frame rate of 100 or 145 Hz (as specified), a camera exposure time of 500 μs, 2×2 camera pixel binning, and in a high-resolution configuration (100× magnification using a UPlanSApo 100×/1.40 oil-immersion objective, Olympus; 50 nm/pixel; 256×256-pixel ROIs). For the experiments in **Fig. 4** imaging was performed with a frame rate of 100 Hz, a camera exposure time of 5 μs, 2×2 camera pixel binning, and in a high-throughput configuration (10× magnification using a UPlanSApo 10×/0.40 NA objective, Olympus, Japan; 500 nm/pixel; 28×28-pixel ROIs). For the experiments in **Fig. 2** the configuration was varied as specified in the configuration chart in **Fig. 2a**. The camera pixel binning was varied between 1×1, 2×2, and 4×4 to achieve various final pixel resolutions. Similarly, the objective was varied between the 10x and 100x objective used in the high-resolution and high-throughput configurations described above. As a result of these differences, the exposure time and illumination source brightness were varied to maintain an overall similar brightness level (as measured by a live histogram of pixel intensities) between configurations with the longest camera exposure time used being 500 μs.

Force on the magnetic beads were measured using variance method, following a correction-free approach.^34^ Briefly, for stationary magnet positions, M-280 beads tethered to a 4-kb DNA were tracked along with a nearby fixed bead at 145 Hz, a camera exposure time of 500 μs, 2×2 camera pixel binning, and in a high-resolution configuration (100× magnification using a UPlanSApo 100×/1.40 oil-immersion objective, Olympus; 50 nm/pixel; 256×256-pixel ROIs). Force was estimated as 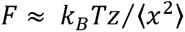 from x-y bead fluctuations measured along the field-parallel direction. This field-parallel direction corresponds to the short-pendulum y-axis in our plots. We confined the force calibration for 4-kb DNA to moderate-force regime to avoid high-force denaturation and pursue force range of interest between 1 to 20 pN. At the upper end of the calibration range (20 pN), the estimated short-pendulum relaxation time is ∼2 ms, with the camera exposure time of 500 μs giving a relaxation-time-to-exposure-time ratio of ∼4.^34^ Under these conditions, residual motion blur may cause a ∼10% overestimation of force.

### DNA substrate construction

The 1 kb DNA containing a 60 nt palindromic hairpin was assembled from three parts. Two separate PCRs were performed on a template plasmid to create the two 500 bp arms. One of the two primers in one of the PCRs contained a 5’ biotin and in the other PCR one primer contained a 5’ digoxigenin. The non-biotinylated DNA end of each PCR product was digested with BsaI (NEB) to create two non-complementary overhangs. An oligonucleotide containing the hairpin sequence and sequences complementary to the two overhangs was then ligated to the two PCR products. This produced a DNA molecule with biotin at one end, digoxigenin at the other, and the hairpin in the middle.

The 4 kb DNA was created from three parts. A plasmid was digested with BsaI (NEB) to create the central 4 kb section with two non-complementary overhangs. Separately two PCRs were performed with the same plasmid as a template to create two 500 bp fragments. Biotin-dUTPs were added to the otherwise standard PCR mix of one of the PCRs. And digoxigenin-dUTPs were added to the other. Thus, one fragment was modified with multiple integrated biotins and the other with multiple digoxigenin. The two fragments were digested with BsaI (NEB) and ligated to the unmodified central 4 kb section creating a DNA with a total length of 5 kb with 500 bp of one end that is randomly interspersed on one end with biotin and the other end with digoxigenin. The resulting 500 bp on each end will be bound to the flowcell or bead surface and not contribute to the extended contour length of ∼1360 nm (4 kb).

### Allan deviation analysis

Allan deviation was calculated from bead-position trajectories using Tweezeypy^25^ Python package. The sampling rate was determined from the median time interval between consecutive frames, and overlapping Allan variance was computed using octave-spaced averaging times. Allan deviation was calculated as the square root of the overlapping Allan variance and reported separately for x, y, and z coordinates. This analysis was used to quantify position stability/resolution as a function of averaging time.

### Hairpin rupture/reformation force

Hairpin rupture and reformation forces were determined from sequential increasing and decreasing force ramps performed on nicked palindromic hairpin DNA. Force was increased from 6 to 20 pN and then decreased at 0.2 pN s⁻¹ while DNA extension was recorded in real time. For each molecule, the rupture or reformation force was defined as the force at which the two extension states were equally populated during the upward or downward ramp, respectively.

### Quality Control (QC) filtering of flow-stretch trajectories

To identify single-DNA tethers and exclude artifact-prone traces, we applied a sequential set of quality-control (QC) filters to bead trajectories obtained from flow-stretching experiments. These filters were designed to remove beads affected by non-specific surface interactions, multiple DNA attachments, unstable baselines, poor tracking quality, or abrupt trajectory discontinuities. Only trajectories that passed all filters were retained for downstream extension analysis.

Filter 1 Return to baseline after flow stops: A subset of beads failed to return to their initial zero-flow position after the flow was stopped, consistent with surface sticking or other non-ideal behavior. We therefore retained only trajectories in which the bead returned to the pre-flow baseline after cessation of flow.

Filter 2 Minimum extension under flow: Partially stuck beads and double- or multi-tethered beads typically exhibited reduced extension under flow relative to singly tethered molecules. To exclude such cases, we retained only trajectories whose flow-induced extension reached at least 50% of the maximum extension observed in that experiment.

Filter 3 Positional noise threshold: To remove poorly tracked or mechanically unstable trajectories, we calculated the standard deviation of bead position during both the zero-flow baseline and the stretched state. Trajectories were excluded if the positional standard deviation exceeded 90 nm during the baseline interval or 200 nm during the stretched interval.

Filter 4 Baseline stability: Stable bead position before flow application was required for reliable extension measurements. We therefore excluded trajectories that exhibited appreciable drift during the zero-flow baseline period (typically 60-90 s), as such drift is indicative of unstable attachment, surface interactions, or tracking artifacts.

Filter 5 Abrupt jump detection: Some trajectories displayed sudden changes in extension during the measurement, likely arising from transient bead-surface interactions followed by release under continued drag. To identify such events, we excluded traces containing abrupt position changes greater than an empirically determined threshold of 800 nm between consecutive samples when the shift persisted for at least 50 frames. In addition, a median absolute deviation (MAD)-based criterion was used to detect extreme discontinuities relative to local trajectory fluctuations.

Together, these filters removed the major classes of non-single-tether and artifact-affected trajectories while preserving beads suitable for quantitative analysis of flow-induced DNA extension.

## Data availability

The data generated in this study are openly available at https://doi.org/10.5281/zenodo.22652860 in Zenodo.

## Code availability

MagTrack and MagScope are available on GitHub at https://github.com/7jameslondon/MagTrack and https://github.com/7jameslondon/MagScope, and stable releases are available through the Python Package Index.

## Acknowledgements

We thank Michael Root of Thomas Jefferson University and Ross Larue of The Ohio State University Wexner Medical Center for their insightful comments and stimulating discussions throughout the preparation of this work. We thank Kathryn Glowinski and Daniel Levine for code review and advice.

## Author Contributions

J.L. and A.S. contributed equally. J.L. wrote the Python code. A.S., J.L., N.T., and T.S. built the magnetic tweezers instrument, performed magnetic tweezers experiments, prepared DNA substrates, and prepared flowcells. J.L., A.S., and N.T. developed the magnetic tweezers experiment protocols. Z.R. ported Python code. The manuscript was written by J.L., A.S., T.S., and R.F. All authors approved the final manuscript.

