## Supplementary Information for "An Accessible Python Framework for Real-Time Magnetic Tweezers Microscope Control and Image Processing"

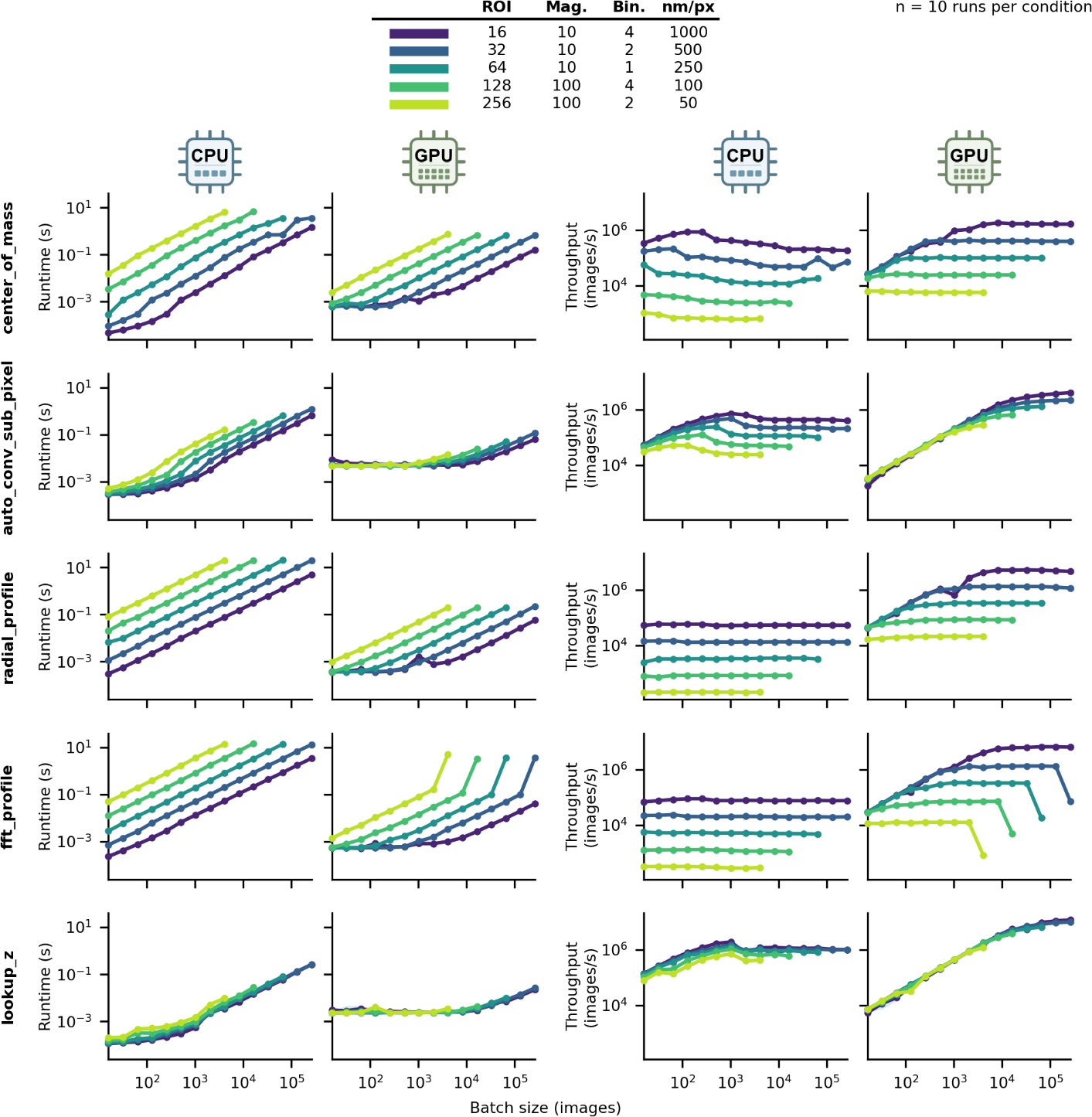

**Supplementary Figure 1. CPU and GPU performance of MagTrack analysis functions.** Runtime and throughput are shown for five MagTrack functions applied to simulated bead-image stacks of increasing batch size. Rows correspond to center of mass, sub-pixel autocorrelation, radial profile, FFT profile, and Z-position lookup; columns show CPU runtime, GPU runtime, CPU throughput, and GPU throughput. Colors denote the ROI, magnification (“Mag.”), pixel binning (“Bin.”), and spatial resolution (“nm/px.”) as specified in the legend at the top of the figure. Points represent the median of 10 runs per condition, with lines connecting batch sizes. Throughput was calculated as batch size divided by runtime. Simulation, memory allocation, CPU-to-GPU transfer, and result validation were excluded from timing.

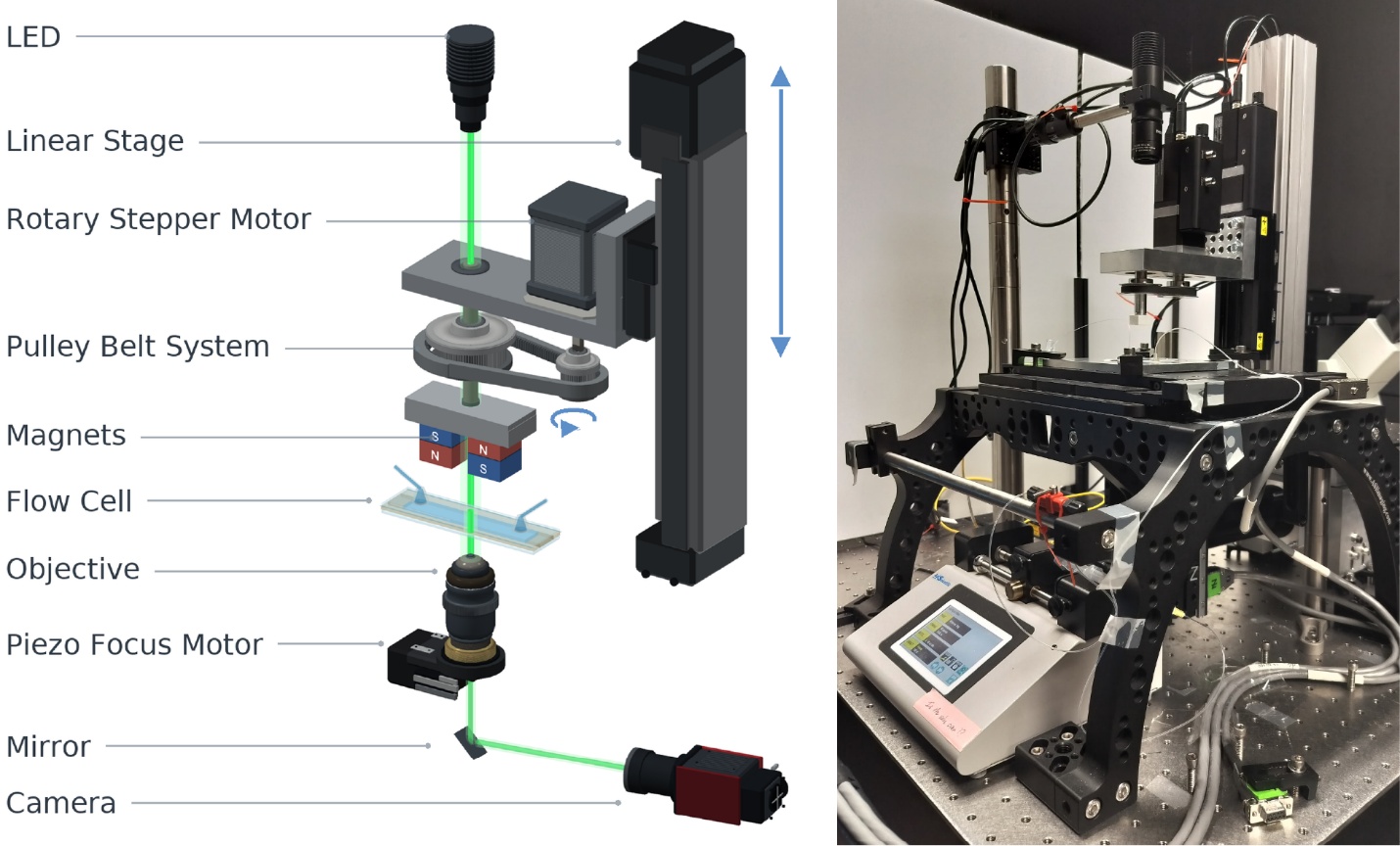

**Supplementary Figure 2. Schematic and image of the magnetic tweezers’ setup.** Left, schematic of the optical and mechanical components of the magnetic-tweezers instrument; right, photograph of the assembled instrument.

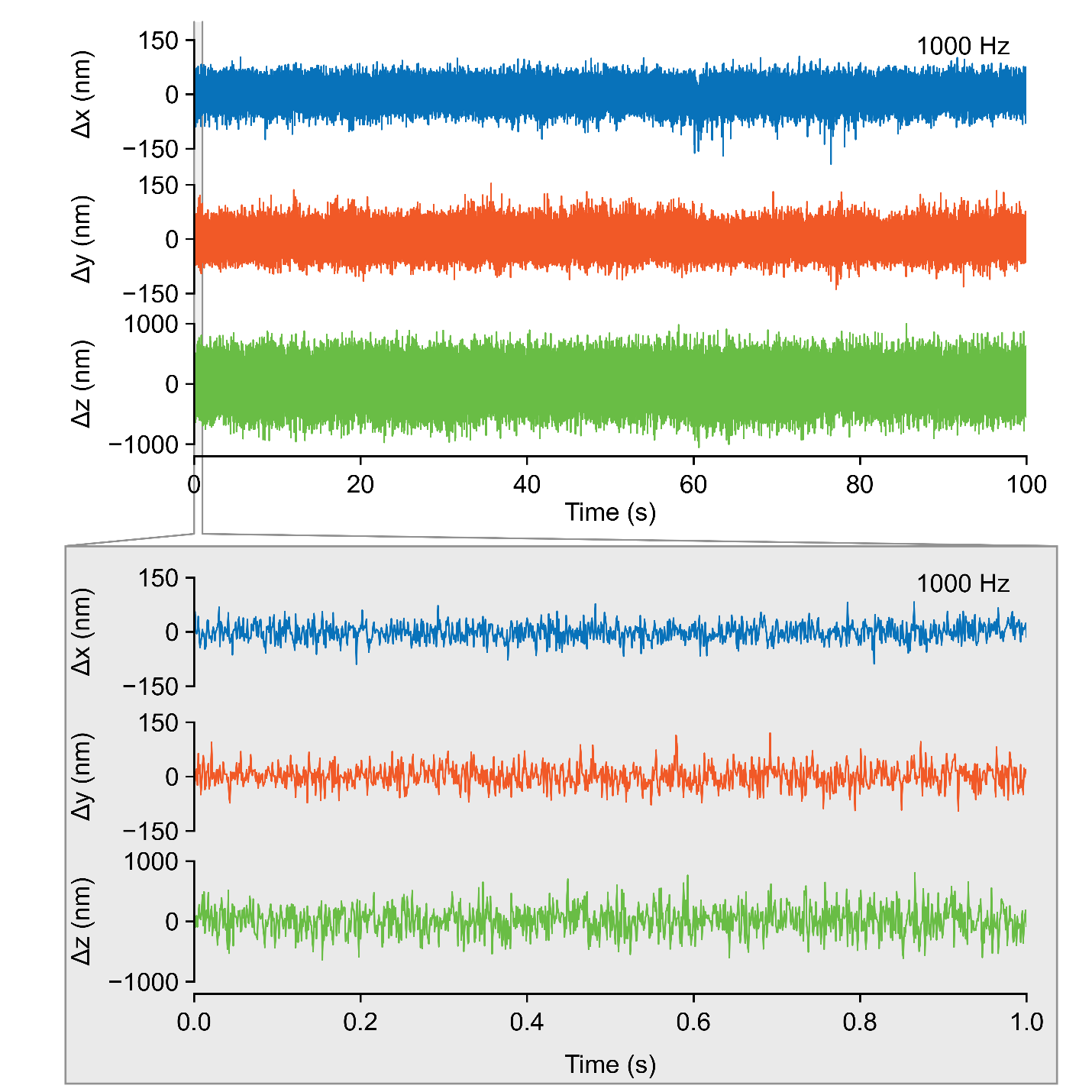

**Supplementary Figure 3. High-rate tracking of fixed reference beads at 1 kHz.** Representative traces of the difference in position between two fixed reference beads acquired at 1 kHz in a high-throughput configuration (10× magnification; 1000 nm/pixel; 16×16-pixel ROIs). Top, 100-s traces; bottom, expanded 1-s interval. On the tested hardware, 237 ROIs were tracked continuously at 1 kHz.

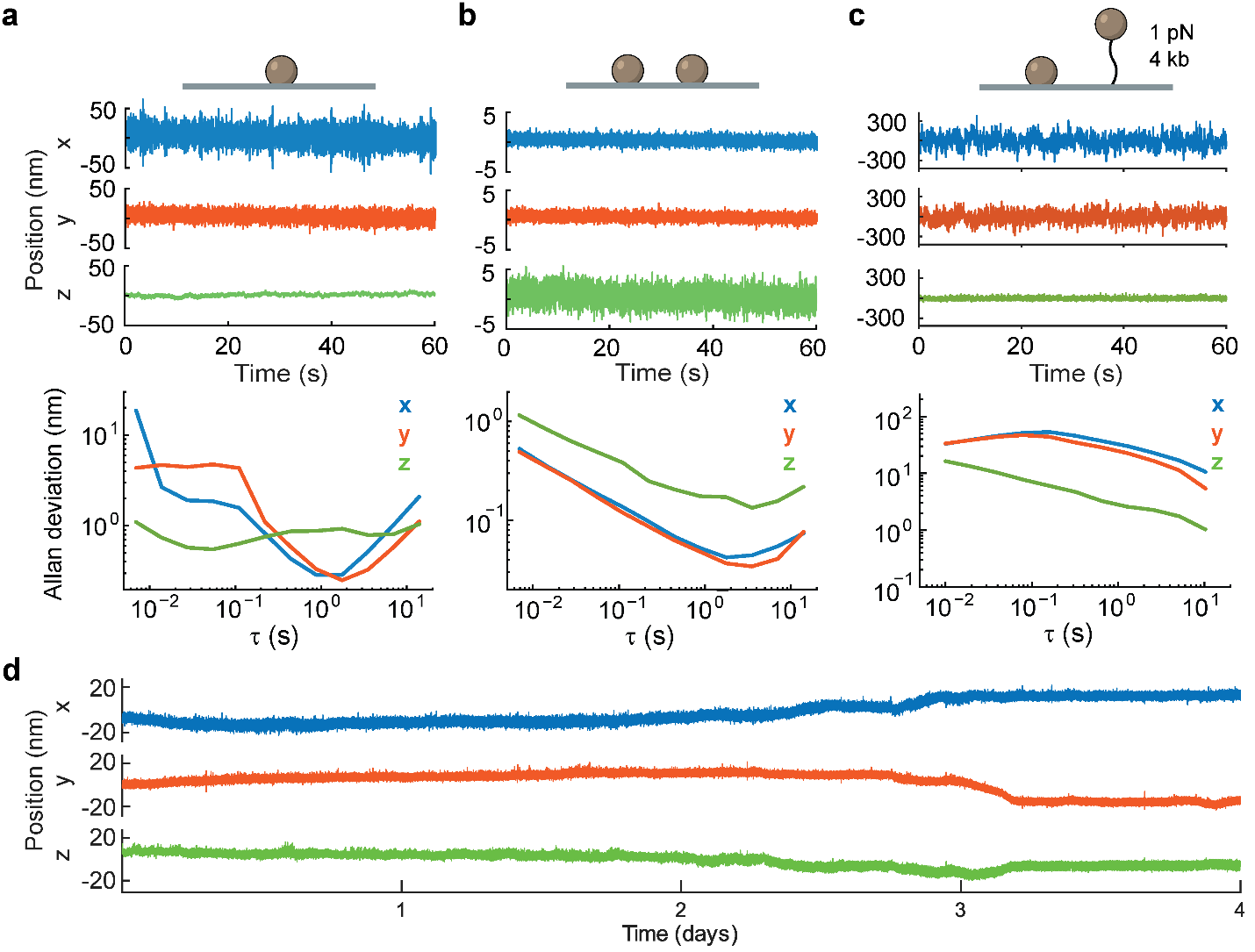

**Supplementary Figure 4. High-resolution fixed-bead and tethered-bead tracking. a** Raw x-y-z position traces and Allan deviations for a fixed 3-μm-diameter polystyrene reference bead tracked at 145 Hz with Z-Lock enabled. **b** Reference-subtracted fixed-bead measurement from the same configuration, obtained by subtracting the position of a second fixed reference bead to remove shared instrument drift and low-frequency vibration. **c** Reference-subtracted tethered-bead measurement from a 2.8-μm streptavidin magnetic bead tethered by 4 kb DNA and held at 1 pN, acquired at 100 Hz; reference subtraction suppresses shared drift while preserving tethered-bead Brownian fluctuations. **d** Extended-duration reference-subtracted fixed-bead x-y-z traces from continuous acquisition lasting more than four days at 100 Hz. All measurements were performed in the high-resolution configuration (100× magnification; 50 nm/pixel; 256×256-pixel ROIs).

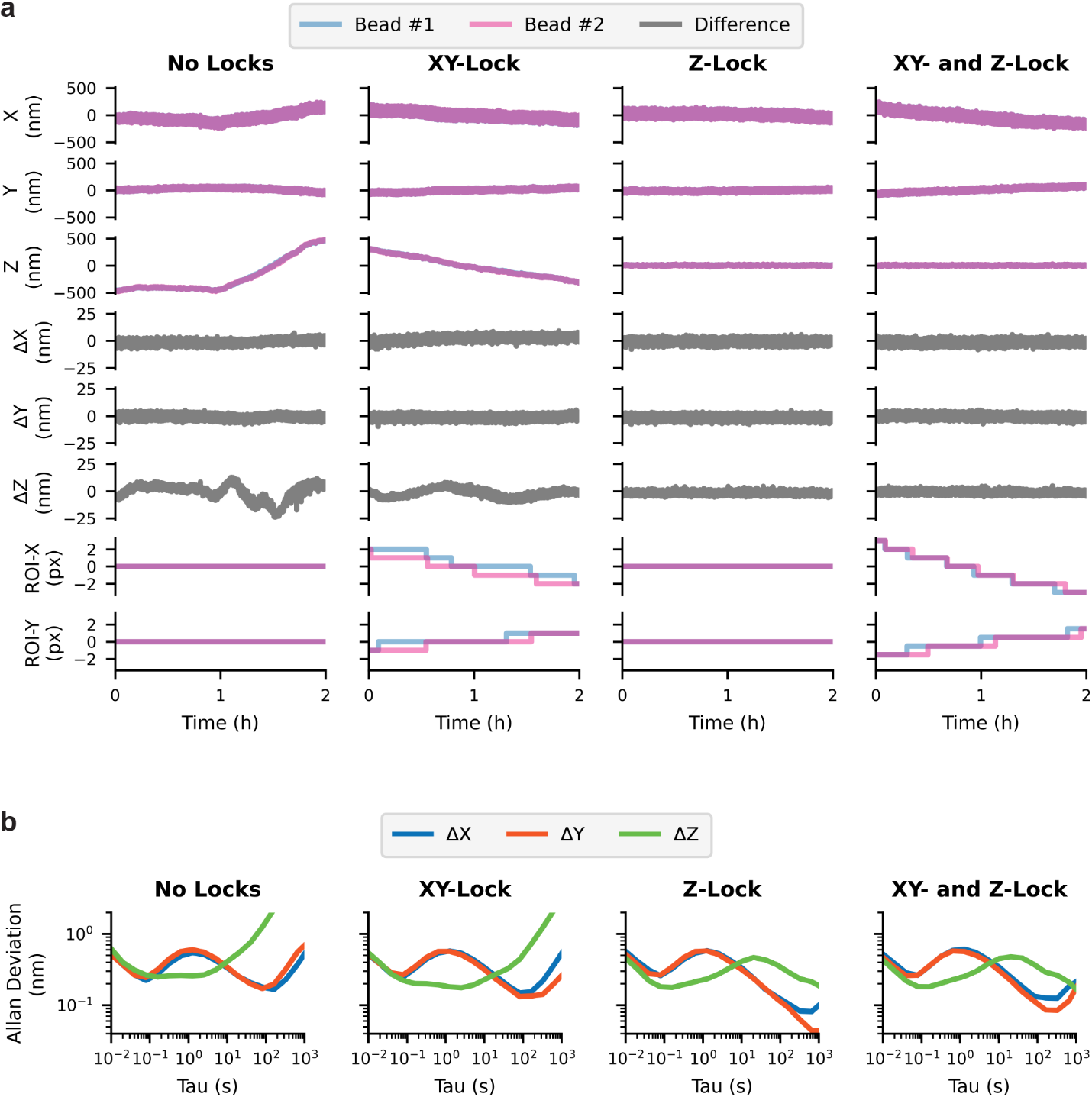

**Supplementary Figure 5. Comparison of tracking with XY-Lock and/or Z-Lock enabled.
a** Raw x-y-z positions, differences between the two bead positions, and x-y ROI coordinates for two fixed 3-μm-diameter reference beads. Measurements were acquired at 100 Hz in a high-resolution configuration (100× magnification; 50 nm/pixel; 256×256-pixel ROIs). **b** Corresponding Allan deviations of the differential traces in panel a.

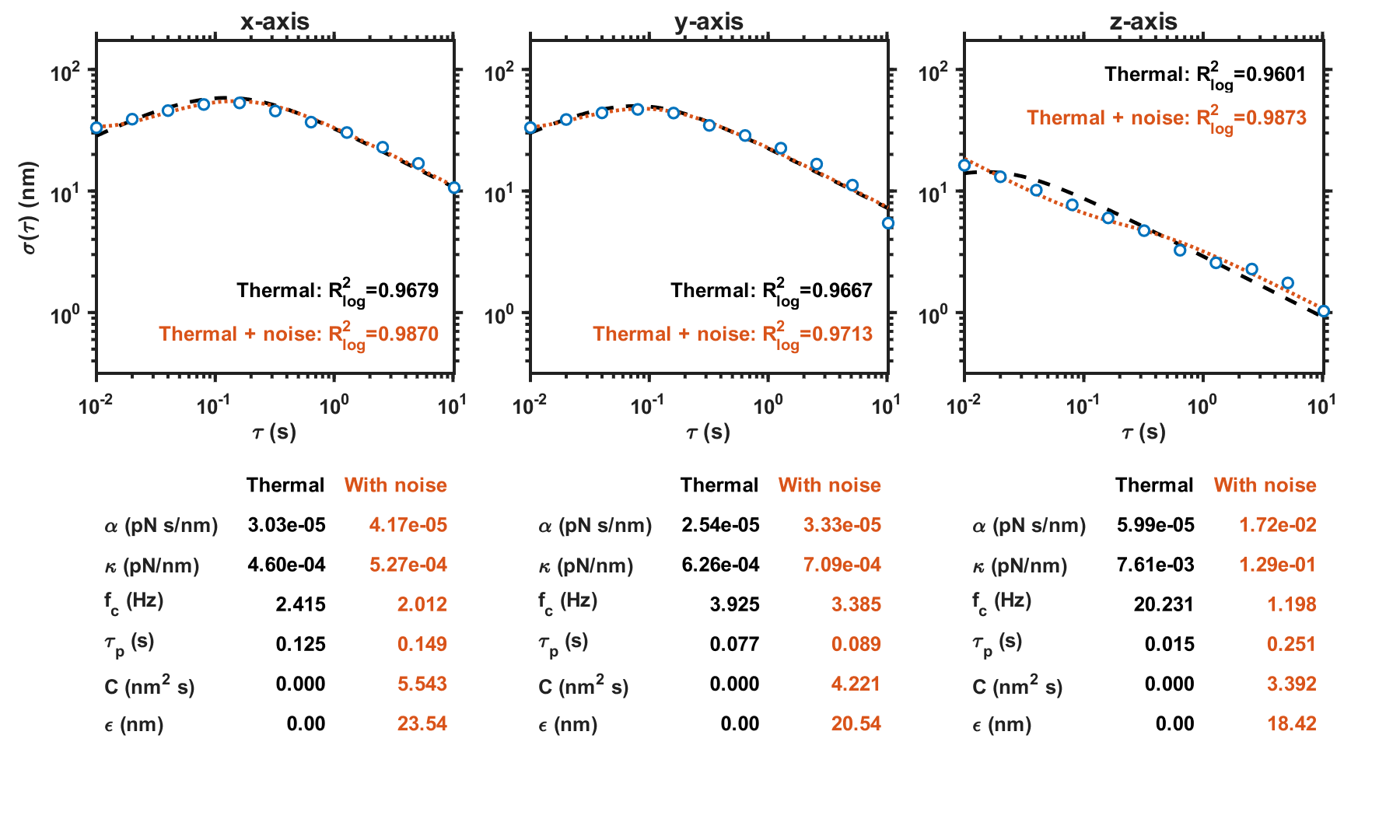

**Supplementary Figure 6**. **Comparison of measured Allan deviations with the analytical prediction for a damped harmonic oscillator.** Allan deviations for the reference-subtracted tethered bead in the x, y and z directions are shown together with curves obtained by fitting to an analytical model of a DNA-tethered magnetic bead in an idealized harmonic trap. Blue circles show measured values; black dashed curves show thermal only fits; orange dotted curves show fits including independent per-frame position noise. The analytical expression is included as a reference to the idealized harmonic-well model underlying single-particle fluctuations in magnetic tweezer experiments. The equation was used in the form,

$\sigma\left( \tau\right)= \sqrt{\frac{2k_{B}T\alpha}{\kappa^{2}\tau} \left( 1+\frac{2\alpha}{\kappa\tau}e^{\frac{-\kappa\tau}{\alpha}}-\frac{\alpha}{2\kappa\tau}e^{\frac{-2\kappa\tau}{\alpha}}- \frac{3\alpha}{2\kappa\tau} \right)+ \frac{C}{\tau}}$,

where $\sigma\left( \tau\right)$ is the Allan deviation, $\alpha$ is the effective drag coefficient, $\kappa$ is the trap stiffness, $k_{B}$ is the Boltzmann constant, and $T$ is the absolute temperature. The term $C/\tau$ represents independent per-frame position noise, with $C= \varepsilon^{2}\Delta t$, where $\varepsilon$ is the effective per-frame noise in the reference subtracted position and $\Delta t$ = 0.01 s is the frame interval (100 Hz). Setting C = 0 gives the thermal-only expression. The displayed $R^{2}$ values and fitted parameters are color-matched to the corresponding curves.

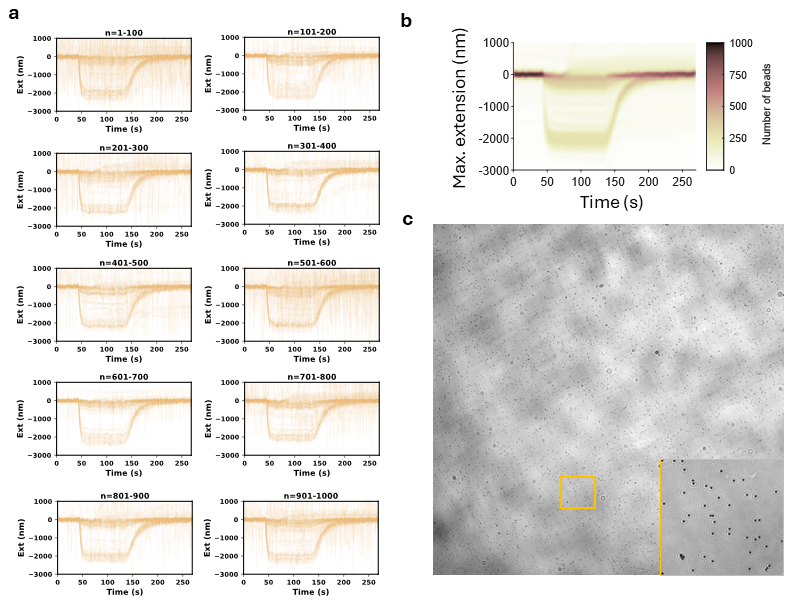

**Supplementary Figure 7. Unfiltered trajectories from 1,000 beads in the flow-stretching assay. a** Raw extension trajectories from 1,000 tracked beads; each of the ten subpanels shows 100 trajectories. **b** Two-dimensional kernel-density map of all overlaid extension trajectories. **c** Representative field of view acquired at 10× magnification; inset, magnified view of the boxed region.
